# DNA opening during transcription initiation by RNA polymerase II in atomic detail

**DOI:** 10.1101/2022.02.05.479244

**Authors:** Jeremy Lapierre, Jochen S. Hub

## Abstract

RNA polymerase II (RNAP II) is a macro-molecular complex that synthesizes RNA by reading the DNA code, a process called transcription. During transcription initiation, RNAP II opens the double-stranded DNA to expose the DNA template to the active site. The molecular interactions driving and controlling the DNA opening are not well understood. We used all-atom molecular dynamics (MD) simulations to obtain a continuous atomistic pathway for the DNA opening process in human RNAP II. To achieve such large-scale and highly nonlinear transition, we steered the MD simulations along a combination of collective variables involving a guided DNA rotation and a set of path collective variables. The simulations reveal extensive interactions of the DNA with three protein loops near the active site, namely the rudder, fork loop 1, and fork loop 2. According to the simulations, these DNA–protein interactions support DNA opening by attacking Watson-Crick hydrogen bonds, and they stabilize the open DNA bubble by the formation of a wide set of DNA–protein salt bridges.

## Introduction

Transcription of DNA to RNA is catalyzed by RNA polymerases (RNAPs), a cornerstone of the central dogma of molecular biology.^1^ In eukaryotes, RNAP II carries out the synthesis of coding RNAs and of many non-coding RNAs. Transcription involves three main steps: initiation, elongation and termination. To trigger initiation, the 12-subunits RNAP II first assembles with general transcription factors to form the pre-initiation complex (PIC).^2^ Within the 12 RNAP II subunits, RNA polymerase subunits 1 and 2 (RPB1 and RPB2, respectively, Fig. 1A) form the cleft and the active site. Several loops protrude from the two large subunits (Fig. 1A), which are well conserved among eukaryotes, including the rudder (in RPB1), fork loop 1 (FL1, in RPB2), and fork loop 2 (FL2, in RPB2).^3,4^ During initiation, these loops are in proximity with the DNA as the transcription bubble forms. The architecture of RNAP II and the mechanism of transcription initiation have been described in several excellent reviews. ^5,6^

**Figure 1:**
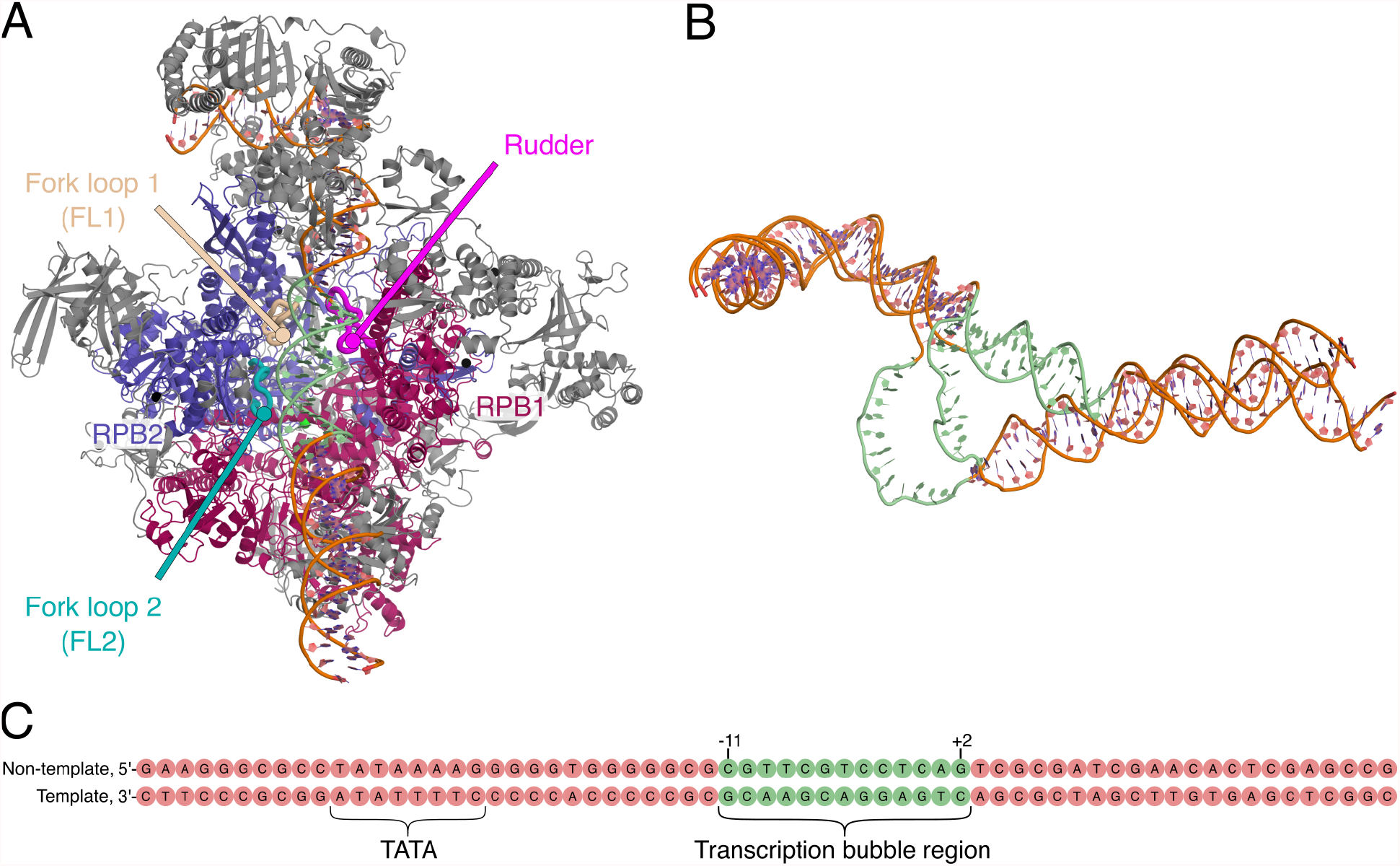
PIC complex in CC and overlap of DNA in CC and OC. (A) Cryo-EM structure of the CC without TFIIH and TFIIS (pdb code 5IY6^9^). Zinc ions shown as black spheres. (B) Overlay of the DNA in CC and OC, taken from structures 5IY6 and 5IYB respectively. ^9^ The DNA region involved in the DNA bubble formation is highlighted in green. (C) DNA sequence simulated in this work, corresponding to the DNA sequence found in 5IYB. DNA numbering according to Ref. 9, where +1 refers to the transcription start site in the OC structure.

Structural studies provided snapshots of the two end states of the PIC during transcription initiation in eukaryotes:^2,7–15^ snapshots of the closed complex (CC), in which DNA is double-stranded and located on top of the RNAP II cleft, and of the open complex (OC), in which the transcription bubble has formed and is loaded into the active site (Fig. 1A and 1B). While a cryo-EM structural study of the bacterial RNAP also revealed intermediate states of DNA opening, ^16^ atomic details of the DNA opening pathway during transcription initiation in eukaryotes are missing. Consequently, the roles of conserved amino acid motifs of the rudder and of FL1 and FL2 during transcription initiation are largely unclear.

Previous molecular dynamics (MD) simulations focused on the elongation step of transcription^17–23^ and on the clamp dynamics during initiation in bacterial RNAP.^24^ A recent coarse-grained MD study addressed DNA melting by inserting DNA base mismatches.^25^ However, DNA opening has not been simulated with atomistic models or without DNA base mismatches.

In this work, we used MD simulations to obtain a continuous opening transition from the CC to the OC in atomic detail. Because the CC-to-OC transition involves conformational rearrangements on the scale of several nanometers, obtaining such transition by brute-force MD simulations is computationally prohibitive. Therefore, we used steered MD simulations^26,27^ along a set of collective variables (CVs) to drive DNA opening and to enhance the sampling along the DNA opening pathway. Our CC-to-OC simulation provides insight into the spatial rearrangements of the DNA and of the protein loops during initiation, and they reveal extensive polar interactions of the DNA with the rudder, FL1, and FL2. These observed interactions suggest roles of the protein loops in supporting DNA strand separation and in stabilizing the transcription bubble in the OC.

## Results

### Steering a 55 °A-conformational transition with a combination of collective variables

Upon forming the transcription bubble, DNA carries out a transition involving a rotation of the DNA double strand by ∼370° as well as a translation of the DNA strands by up to 55 °A relative to the protein.^9^ Simulating such large-scale, nonlinear conformational transitions in atomic detail imposes considerable challenges. One possible strategy for favoring these large-scale motions is to introduce base mismatches between the two DNA strands, as used for obtaining the OC cryo-EM structure by He *et al*.^9^ or used for favoring DNA melting in coarse-grained MD simulations.^25^ In contrast to these previous studies, we simulated DNA opening without base mismatches, according to the biological relevant state of the CC (Fig. 1C). We obtained a relaxed pathway of DNA opening with a combination of two methods. First, we obtained an initial pathway using steered MD simulations along a combination of three collective variables (CVs); second, the initial pathway was relaxed using the path collective variable (PCV) method.^28^

To guide the opening pathway, steered MD simulations were carried out along a combination of the following three CVs: (i) a rotational CV applied to the downstream DNA helix, thereby driving the melting of the DNA strand (Fig. 2A, *ξ*_1_); (ii) two CVs given by the root mean-square distance (RMSD) of the sugar-phosphate backbone of the DNA relative to the conformation in the OC, taken from the 5IYB structure (*ξ*_2_ and *ξ*_3_).^9^ Figure 2A illustrates the evolution of the rotational CV *ξ*_1_ and of the RMSD-based CV *ξ*_3_. By pulling along these CVs we obtained an initial path of DNA opening (Movies S1 and S2).

**Figure 2:**
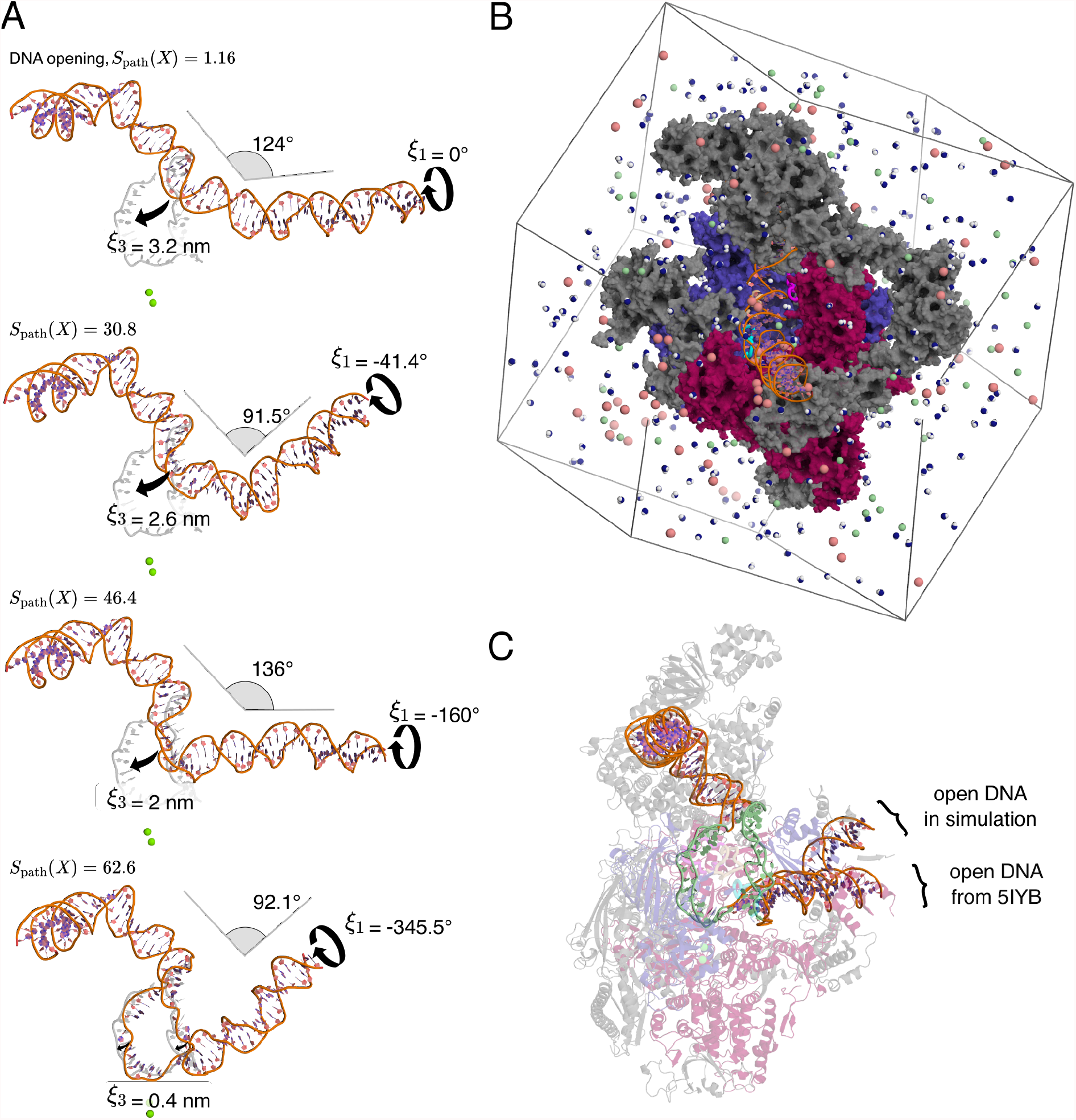
Transition from closed to open DNA in atomic detail. (A) Opening transition snapshots with corresponding *S*_path_ value (progression along the DNA opening), rotation of the downstream DNA helix (*ξ*_1_), RMSD relative to the open bubble (*ξ*_3_), and DNA helix bending angle. The target open bubble conformation is depicted in light gray. For reference, two catalytic magnesium ions are shown as green spheres. (B) Simulation box. Colors for the PIC in CC are consistent with Fig. 1. Water molecules, sodium ions, and chloride ions are colored in blue and white, pale pink, and pale green respectively. Most of water molecules and ions have been removed for clarity. (C) Section of the PIC in OC obtained from steered MD simulations. Open DNA from 5IYB structure is shown for reference. The transcription bubble is colored in pale green.

### A path collective variable (PCV) for steering and relaxing the DNA opening path

Because our steered MD simulations were carried out on much shorter time scales compared to experimental time scales, it is reasonable to believe that the initial path is still biased by non-equilibrium effects. To relax the conformations along the opening pathway and, thereby, to mitigate such non-equilibrium effects, we applied the PCV method.^28^ Generally, PCVs are defined using two CVs: the position *S*_path_ along the initial path and the distance *Z*_path_ from the path, where the path is defined along *N* intermediate conformations (see Methods). In this study, the initial PCV was defined with 72 intermediate conformations taken form the steered MD simulation. Then, we carried out two rounds of constant-velocity pulling along *S*_path_. Within each round, the path was allowed to relax, providing us with an updated set of increasingly relaxed intermediate conformation and, thereby, an updated PCV. The final PCV along 63 relaxed intermediate conformations, allows convenient opening simulations by pulling along the single *S*_path_, instead of pulling along the three CVs used for obtaining the initial path (see above). In addition, projection onto the final *S*_path_ provides a convenient measure for the progress of the opening pathway, as used below in our figures and analysis. More details are provided in the Methods.

### Atomistic transition from the closed to a stable open DNA

By pulling along the aforementioned *S*_path_, we obtained all-atom continuous trajectories of DNA opening from the CC to the OC (Fig. 2A/C and Movies S1, S2). To test whether we have reached a state with a stable open DNA bubble, we simulated the final state without any biasing potential for 200 ns. In this simulation, the distances between the disrupted base pairs were reasonably stable (Fig. S1A, B), demonstrating that the strands did not re-anneal, as expected for a stable OC. The open DNA bubble exhibited a length of 12 base pairs (bp), in reasonable agreement with the length of 13 bp in the reference structure by He *et al*.^9^ We quantified the spatial extension of the bubble with the average distance *d*_bb_ between the 12 disrupted base pairs. In the unbiased 200 ns simulation of the OC, we obtained *d*_bb_ = 2.36 nm (Fig. S1), in reasonable agreement with the value of 2.67 nm in the reference structure. Minor structural differences relative to the reference OC are expected because (i) the DNA bubble is flexible and (ii) RNAP II in the OC accommodates various DNA bubble lengths and widths during initiation and, more generally, during the entire transcription process.^29^ Overall, the stability of the DNA transcription bubble in our free simulation implies that the previous pulling simulation represents a complete DNA opening pathway.

Figure 2A and Movies S1 and S2 provide an atomic view on the large DNA rearrangements. First, due to the clockwise rotational motion carried out by the downstream DNA, the DNA became underwound in the transcription bubble region. Second, due to the translational motions induced on the transcription bubble towards the active site, the DNA bent at the transcription bubble region. These two topological changes of DNA led ultimately to the disruption of 12 bp. DNA rotational angles, bending angles and RMSD relative to the reference OC are depicted in Fig. 2A for four snapshots of our DNA opening trajectory. This interplay between negative supercoiling (clockwise rotational motion of DNA), bent DNA and base pair disruptions have been reported previously in DNA minicircles.^30–33^ In addition, negative supercoiling has been shown to promote DNA opening during transcription with minimal transcription factors, ^34^ further corroborating that our simulations reflect experimentally relevant conditions. Together, we obtained a simulation protocol that provides continuous atomistic transitions from the CC to a stable OC with an open DNA transcription bubble within computationally accessible simulation times. The protein–DNA contacts and interaction energies obtained from the simulations are discussed in the following sections.

### Fork loop 2 tilting during DNA opening

Because DNA opening occurs inside the PIC, DNA extensively interacts with protein domains and, in particular, with the protein loops. While DNA was loaded into the active site in the simulations, FL2 tilted into the transcription bubble, in-between the two DNA strands (Fig. 3A-D and Movie S3). Whereas solvent-exposed protein loops are often flexible, the FL2 conformation pointing into the open bubble was remarkably stable, locked by electrostatic protein–DNA interactions, as observed in the free 200 ns simulation following the opening transition described above. The FL2 tilting in our simulations is compatible with a hypothesized role of FL2 as a sensor for the open transcription bubble.^9^ A recent study^15^ revealed a similar conformational change of FL2 during the transition from the CC to the OC, further supporting that FL2 is acting as a sensor for DNA opening.

**Figure 3:**
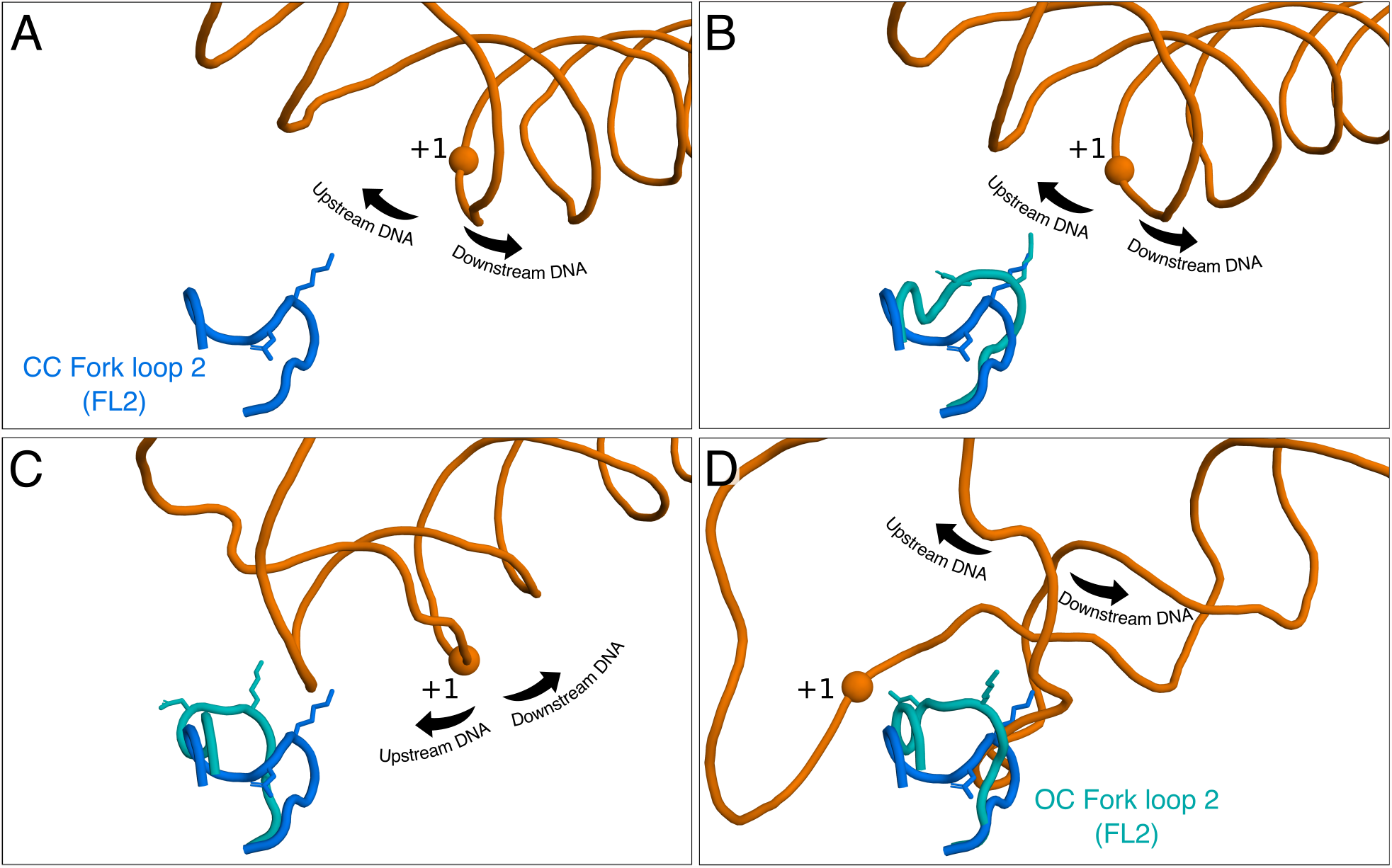
Tilting of sensor fork loop 2 (FL2) into the transcription bubble during DNA opening. (A-D) Snapshots of FL2 (cyan) during DNA opening corresponding to *S*_path_ = 1.16, 31.8, 51.5 and 62.6 respectively. The final tilted state of FL2 is depicted in panel D. For reference, the starting position of FL2 (marine blue), Asp-492 and Lys-494 are shown.

### Fork loops 1 and 2 support DNA opening by hydrogen bond attack on Watson-Crick pairs

The simulations revealed how DNA opening is supported by the rearrangement of hydrogen bonds (H-bonds) between DNA, protein, and water, as shown in Fig. 4. Namely, the loss of 31 Watson-Crick (WC) DNA–DNA H-bonds (Fig. 4A, orange curve) was predominantly compensated by the formation of approximately 40 DNA–Water H-Bonds (Fig. 4A, blue curve). The open DNA was further stabilized by the formation of approx. 12 DNA–protein H-bonds (Fig. 4A, green curve), among which ∼50% formed with the Watson-Crick edge (Fig. 4A, red curve), and the other ∼50% formed with other edges or with the DNA backbone.

**Figure 4:**
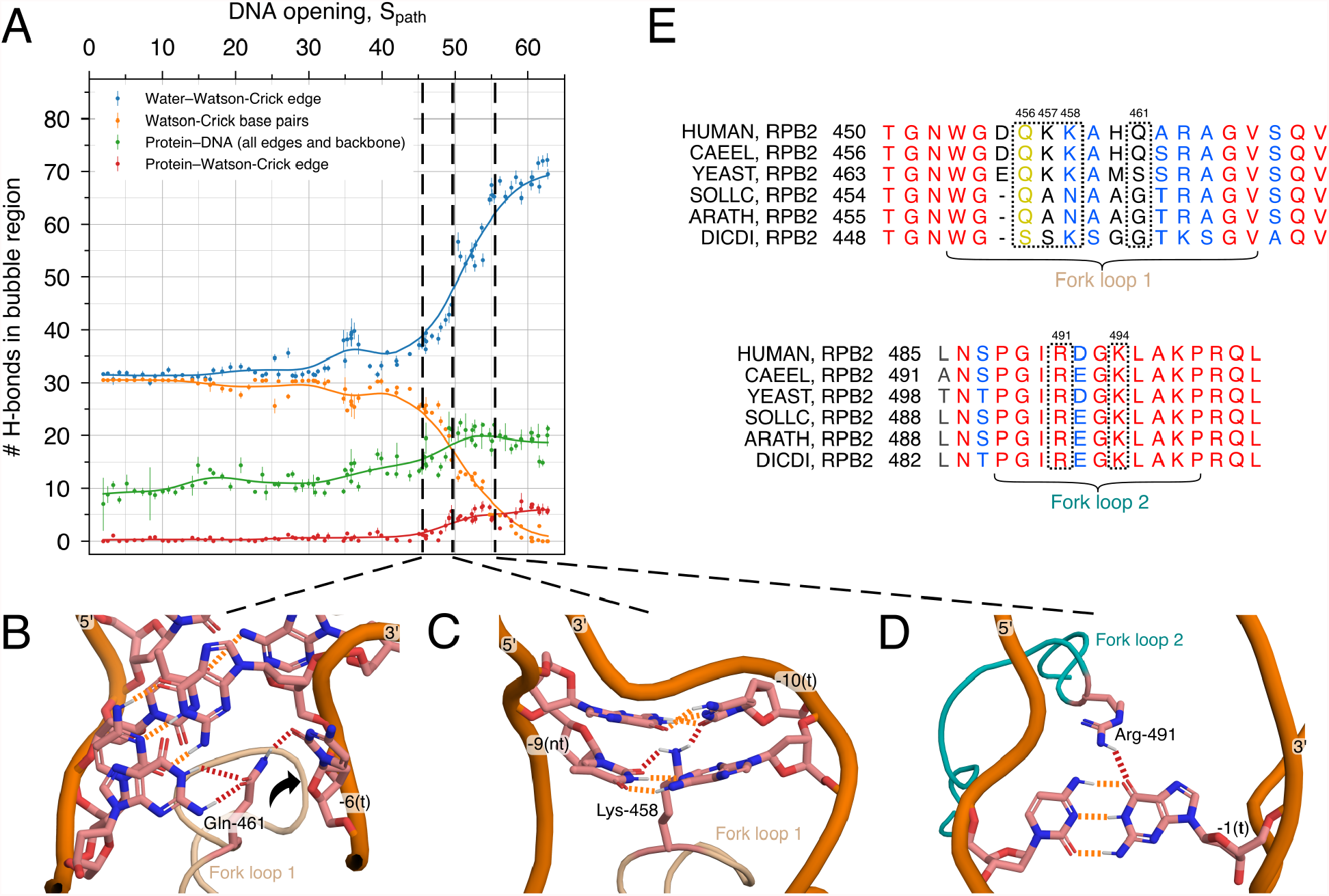
Rupture of Watson-Crick H-bonds in the transcription bubble and formation of DNA–protein and DNA–water H-bonds during the DNA opening process. (A) Development of the H-bonds of the transcription bubble region: number of H-bonds between WC edge and water (blue), between base pairs (orange), between DNA and protein (green), and between DNA WC edge and protein (red). Smooth lines are shown to guide the eye. (B) WC H-bond disruption driven by Gln-461 and base flipping (black arrow) of DNA residue 6 in the template strand. (C–D) Attack of WC H-bond by fork loops 1 and 2, respectively. (E) Sequence alignment (CLUSTAL W^35^) of fork loop 1 and 2 from six different eukaryotes: *Homo sapiens, Caenorhabditis elegans, Saccharomyces cerevisiae, Solanum lycopersicum, Arabidopsis thaliana* and *Dictyostelium discoideum*. The residue color highlights the degree of conservation: invariant residues (red), residues with similar properties (blue), or with weakly similar properties (yellow). Residues in dotted boxes are discussed in the text.

The progression of WC H-bonds (Fig. 4A, orange curve), together with visual inspection of the MD trajectories, revealed three key protein residues involved in destabilizing the double-stranded DNA by attacking the Watson-Crick H-bonds. These events are reflected by marked decreases of the number of WC H-bonds at *S*_path_ = 45.9, 49.6, and 55.4 (Fig. 4A, vertical lines) and are visualized in the molecular representations of Figs. 4B–D. First, the side chain of Gln-461 of FL1 interacted with the base pairs at −6, thereby competing with the WC base pairing (Fig. 4B, *S*_path_ = 45.9). Second, the side chain of Lys-458 of FL1 interacted via H-bonds with base pairs at position −9 and −10 (Fig. 4C, *S*_path_ = 49.6). Third, at a later stage of the opening process and after FL2 tilted into the open bubble, Arg-491 of FL2 destabilizes WC H-bonds, thereby promoting the unzipping of the double-stranded DNA (Fig. 4D, *S*_path_ = 55.4). Hence, DNA–protein interactions do not merely serve as a compensation for the loss of DNA–DNA interactions in order to energetically stabilize the final OC state, but they might also catalyze the rupture of the WC base pairing.

The DNA dynamics in RNAP II driven by FL1 and FL2 are not unique but instead resemble dynamics observed in other DNA-interacting enzymes. For instance, base flipping has been suggested as an early mechanistic stage for DNA opening in a bacterial promoter.^36^ Likewise, H-bond attack to WC base pairs has been proposed for the cytosine 5-methyltransferase, where the enzyme infiltrates the DNA helix by forming H-bonds with nucleic bases, consequently destabilizing WC H-bonds and inducing base flipping.^37,38^ Similarly, a base flipping event at position −6 of the template strand occurred during DNA opening in our simulation. Here, base flipping was promoted by the aforementioned Gln-461, via disruption of the WC H-bonds during the DNA opening (Fig. 4B, black arrow).

To get additional insights into the role of DNA–Protein interactions during DNA opening, and to identify selection pressure on the three key residues mentioned above, we analyzed the residue conservation of FL1 and FL2 among six eukaryotic organisms by means of multiple sequence alignments. Overall, FL1 and FL2 are strongly conserved among eukaryotes demonstrating their critical biological roles (Fig. 4E). However, Gln-461 is not conserved among eukaryotes, suggesting that the DNA–Gln-461 interactions observed in our simulations is either not critical for DNA opening or may be replaced with other interactions. In contrast, Lys-458 is well conserved among the analyzed eukaryotes (Fig. 4E); we hypothesize that the substitutions with Asn in *Solanum lycopersicum* and *Arabidopsis thaliana* may interact with DNA similar to Lys, thus supporting the role of residue 458 in destabilizing the double-stranded DNA. Arg-491 is invariant among all the eukaryotic organisms chosen here (Fig. 4E), underlining its biological relevance in DNA strand separation. Taken together, our data suggest that residues of FL1 and FL2 catalyze DNA opening by attacking WC H-bonds between double-stranded DNA, providing a rationale for the marked sequence conservation of FL1 and FL2. According to the simulations and the sequence alignment, the conserved residues Lys-458 and Arg-491 are involved in H-bond attack; however, we cannot exclude the possibility that other conserved residues play similar roles.

### Fork loops–DNA and rudder–DNA electrostatic interactions stabilize the open DNA conformation

To rationalize the energetic driving forces for DNA opening, we monitored the potential energy from DNA–DNA, DNA–Protein, and DNA–Water interactions (Fig. 5A). Here, potential energies were taken as the sum of Lennard-Jones and short-range Coulomb interactions, averaged over 50 ns of simulation and normalized relative to the state of the CC. The loss of interactions between the DNA strands is primarily compensated by a large gain of DNA–Protein interactions, as evident from the large negative DNA–Protein potential energies (Fig. 5A, orange and red). Although DNA opening leads to an increase of DNA– water interactions, as expected from the formation of H-bonds between water and the WC edge (Fig. 4A, blue), DNA–water interactions (Fig. 5A, blue) play a much smaller role as compared to DNA–Protein interactions (Fig. 5A, red).

**Figure 5:**
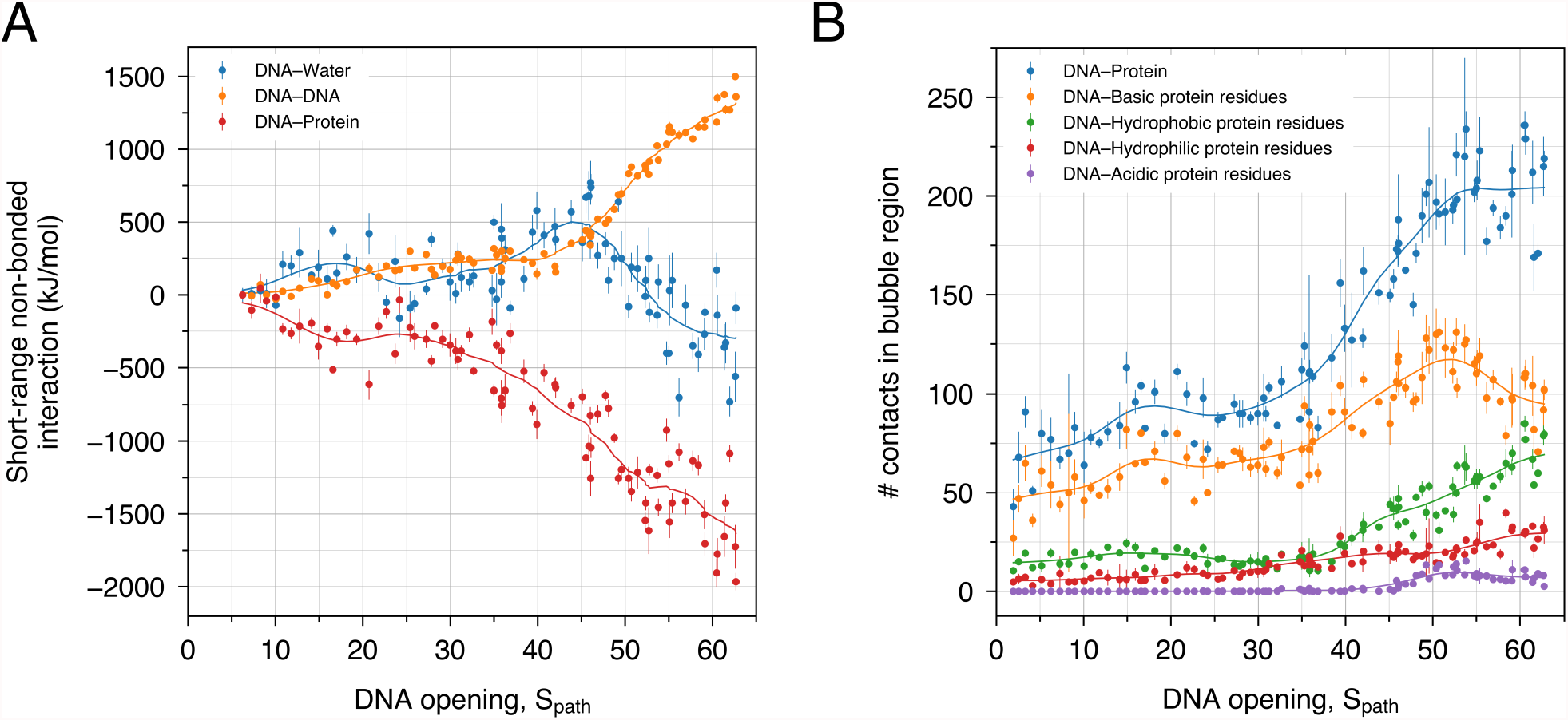
Electrostatic interactions support DNA opening. (A) Coulomb and Lennard-Jones short-ranged interactions between DNA and water (blue), between DNA and DNA (orange) and between DNA and protein (red). (B) Number of contacts within the DNA bubble region with all protein residue types (blue), with basic protein residues (orange), with hydrophobic protein residues (green), with hydrophilic protein residues (red) and with acidic protein residues (purple). Smooth lines are shown to guide the eye.

To quantify which type of DNA–Protein interactions drive DNA opening, we further analyzed the number of contacts of the DNA bubble region with different groups of amino acids of common physicochemical properties (Fig. 5B). Evidently, the DNA forms ∼50 new contacts with basic protein residues, far more compared to contacts with polar or acidic residues. This finding reflects that RNAP II cleft is highly positively charged which helps to attract the negatively charged DNA backbone deeper into the cleft and, in particular, into the active site. This finding demonstrates, not surprisingly, that electrostatic interactions between the DNA and RNAP II are the key energetic driver for transcription bubble formation.

Visual inspections of the simulations revealed reoccurring salt bridges and hydrogen bonds between the protein and the open DNA. Gln-456 and Lys-457 (in FL1) form H-bonds with the template strand of DNA (Fig. 6A), suggesting that FL1 stabilizes the open bubble by compensating the loss of H-bonds between the two DNA strands and by imposing a steric obstacle against strands re-annealing. In close proximity to FL1, Arg-327 (in the rudder) forms a salt bridge with the template strand (Fig. 6B). Likewise, Lys-494 (in FL2) and Arg-222 (in RPB2) interact with the non-template strand with salt bridges and H-bonds (Fig. 6C). Arg-222, Lys-413 and Arg-416 are examples of residues outside the fork loops or the rudder that form electrostatic interactions with the non-template strand of the open DNA conformation (Fig. 6D).

**Figure 6:**
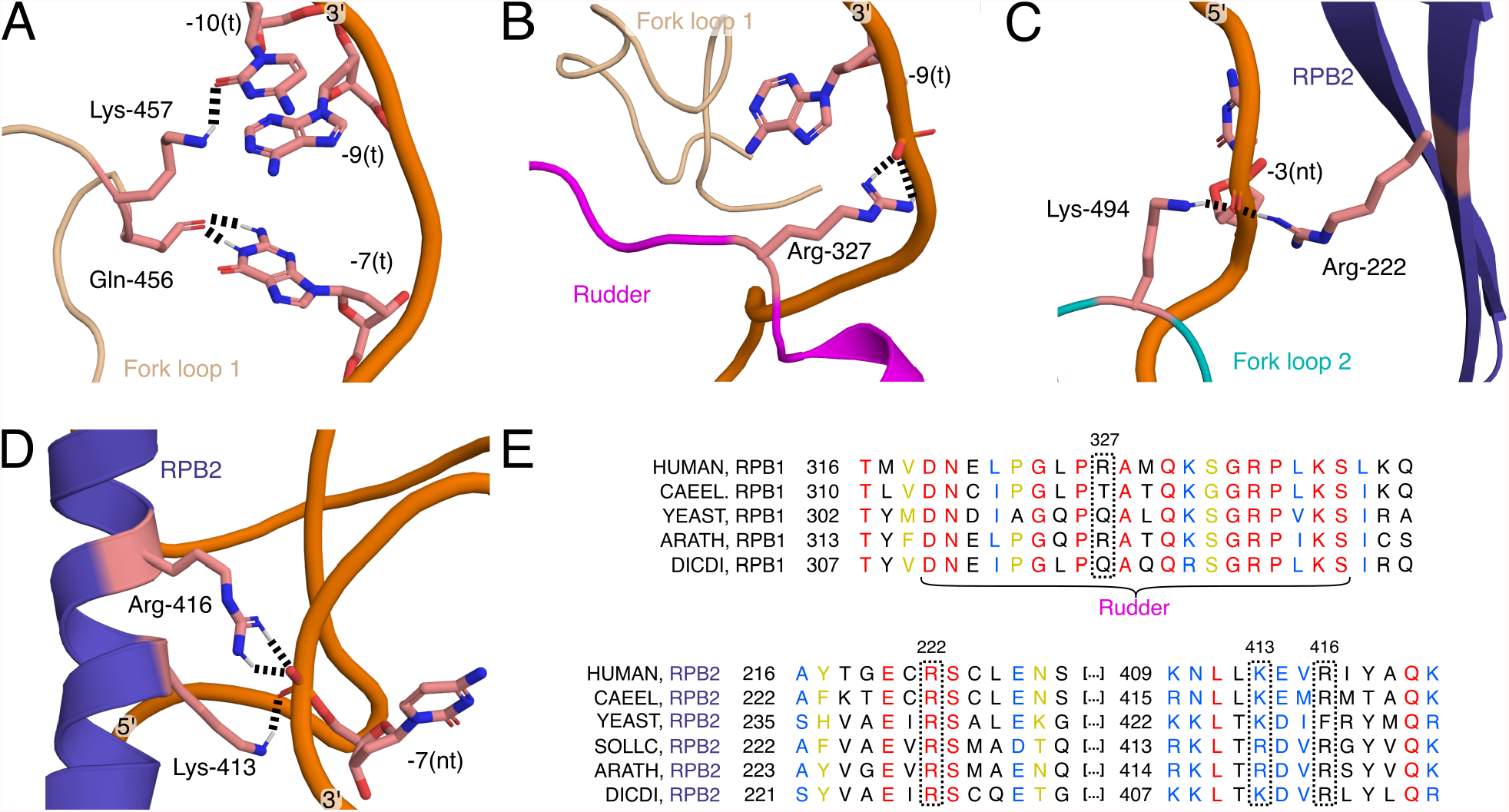
Electrostatic interactions between DNA and PIC stabilize open DNA in the OC. (A-D) H-bonds between protein and DNA. (B-D) Salt bridges between cationic residues with the anionic DNA backbone. (E) Sequence alignment (CLUSTAL W) of the rudder from different eukaryote organisms. Red residues are invariant, blue are from groups of strongly similar property and yellow from groups of weakly similar properties.

To corroborate the relevance of the DNA–protein interactions observed in our simulations for stabilizing the transcription bubble, we inspected the conservation of the residues mentioned above with sequence alignments. Accordingly, Lys-494 (Fig. 4E, FL2) and Arg-222 (Fig. 6E, RPB2) are invariant while Lys-413 (Fig. 6E, RPB2) is well conserved, supporting their biological relevance. Gln-456 (Fig. 4E, FL1) and Arg-416 (Fig. 6E, RPB2) are largely conserved except in *Dictyostelium discoideum* and in *Saccharomyces cerevisiae* respectively, underlining their putative role in stabilizing the transcription bubble. In contrast Arg-327 (Fig. 6E, rudder) is not conserved but may be replaced with Thr or Gln; however, all those residues are capable of forming H-bonds with the DNA backbone and, thereby may all stabilize the open bubble. Finally, Lys-457 (Fig. 4E, FL1) is not conserved suggesting that this residue is less critical for stabilizing the transcription bubble. Together, our simulations show extensive interactions between the PIC and DNA that stabilize the DNA bubble. In the light of the sequence alignments, many of these interactions are critical, whereas some may be replaced by other interaction.

## Discussion

We have presented the first all-atom simulation of a continuous DNA opening transition within human RNAP II. The simulations revealed extensive electrostatic and polar interactions of the DNA with the protein, predominantly with the two fork loops and with the rudder. Closer inspection of these interactions suggested that the rudder and the two fork loops are involved in (i) the separation of the two DNA strands by means of H-bond attack to WC base pairs and (ii) in maintaining the open DNA conformation by a combination of steric hindrance and electrostatic interactions. The biological relevance of the protein residues involved in the observed interactions was further scrutinized by analyzing their conservation among eukaryotic amino acid sequences. Finally, we observed a base flipping event as well as the flipping of FL2 into the transcription bubble, in line with previous experiments.^9,15,36–38^

Mutagenesis experiments targeting the fork loops and the rudder have been carried out in archaeal RNAP II and have revealed that the rudder helps in stabilizing the melted DNA in the OC.^39^ In addition, these experiments suggested that FL2 and, in particular Arg-451 —the archaeal equivalent of Arg-491 mentioned in this work—, plays a role in unwinding downstream DNA during elongation. However, mutagenesis of FL1 did not impact DNA opening in archaeal RNAP II in this study. Whereas archaeal and human RNAP II exhibit high sequence conservation, the physiological temperature in which DNA opening occurs may strongly differ, which might influence DNA melting. Indeed, a temperature of 70°C was used in the permanganate footprinting experiment by Naji *et al*. compared to a temperature of 37°C expected for human physiological conditions. ^39^ The same kind of mutagenesis experiments in eukaryotic RNAP II system, ideally for human RNAP II, would be highly interesting to confirm that FL1, FL2, and the rudder are essential for DNA opening.

Simulating and relaxing such a large-scale roto-translational conformational transition with atomic MD force fields is computationally challenging. A recent coarse-grained MD study introduced base pair mismatches in the transcription bubble region to favor DNA opening. ^25^ However, if DNA melting occurs without simultaneous rotation of the downstream DNA, high DNA strains in the upstream or downstream DNA region emerge, which is incompatible with the open DNA conformation from experimental structures. ^8,9,40^ Therefore, in this work, we used steered MD simulations along a combination of three CVs to guide the large-scale displacement of DNA by up to 55 °A simultaneously with DNA rotation by ∼ 346°. Henceforth, we used the PCV framework to relax the initial opening pathway from steered MD. Notably, we tried to compute the potential of mean force along the PCV with umbrella sampling^41^ or metadynamics,^42^ with the aim to obtain an estimate for the free energy of DNA opening; however, we observed considerable hysteresis problems, suggesting that it is difficult to sample all the degrees of freedom orthogonal to the PCV. This observation further implies that our simulations provided a plausible pathway for DNA opening, but not necessarily the minimum free energy pathway. Future simulations may investigate the use of additional enhanced sampling techniques such as bias-exchange umbrella sampling ^43^ or the use of extensive compute power. ^44^

In eukaryotes, the transcription factor TFIIH is involved in, both, translocase activity and DNA opening. ^45–48^ However, it has been suggested that translocase activity is not necessary for RNA transcription^49^ and that TFIIH-independent DNA opening is possible in yeast RNAP II^12^ or in human RNAP II under negative supercoiling conditions. ^34^ In this study, we used rotational and translational CVs to model the motions induced by negative supercoiling in TFIIH-independent DNA opening. However, since TFIIH also produces torsional stress to the downstream DNA, our current protocol for DNA opening will be useful to study DNA opening in presence of TFIIH. Simulations with TFIIH will be particularly relevant to understand the role of its XBP subunit, the TFIIH subunit containing the translocase activity and the motor for DNA unwinding. ^47^

The transcription factor TFIIB contains the B-reader and B-linker elements, which also help DNA opening.^10,50^ Because refined atomic models of the B-reader and B-linker were not resolved in the CC structure by He *et al*.,^9^ we simulated the CC-to-OC transition in the absence of these TFIIB elements. Further atomistic simulation will be needed to investigate whether the B-reader and B-linker support DNA opening using similar interaction motifs as observed here for FL1, FL2, and for the rudder.

To conclude, we obtained an atomic model for a continuous DNA opening event in the human PIC. The simulations revealed extensive interactions of the DNA with the protein, in particular with loops protruding into the polymerase cleft: FL1, FL2, and the rudder. According to the simulations, the loops play multiple roles for DNA opening: (i) by attacking WC H-bonds, they may catalyze the melting of the DNA; (ii) extensive polar interactions via salt-bridges with the DNA backbone and, to a lower degree, via H-bonds with the DNA backbone and bases stabilize the open DNA conformation; (iii) FL2 tilted into the DNA bubble during opening, a conformational transition that is compatible with a function of FL2 as a sensor for an open transcription bubble.

## Methods

### Simulation setup

MD simulations were carried out with Gromacs, ^51^ version 2020.2 patched with Plumed ^52,53^ version 2.6.1, and with Gromacs version 2021 patched with Plumed version 2.7.0. The initial atomic coordinates for the CC were obtained from the Protein Data Bank (accession code 5IY6^9^) from which we removed TFIIH and TFIIS. We used YASARA^54^ version 20.8.23 to add acetyl and N-methyl amide capping groups at the ends of the missing protein regions and at the C and N termini. We also used YASARA^54^ to add missing atoms. The system was solvated with TIP3P water molecules^55^ and Na/Cl counter ions were added to neutralize the system with a salt concentration of 100 mM. In total, the system contained 832078 atoms. The OL15^56^ force field was used for the DNA. The ff14sb^57^ force field was used for the protein except for the zinc(II)-coordinating Cis and His residues, for which the improved parameters by Macchiagodena *et al*.^58^ were used.

Electrostatic interactions were computed with the particle-mesh Ewald method, ^59^ using a real-space cutoff at 1 nm and a Fourier spacing of 0.16 nm. Dispersion interactions and short-range repulsion were described with a Lennard-Jones potential with a cutoff at 1 nm. Bonds and angles of water were constrained with the SETTLE algorithm^60^ and bonds involving other hydrogen atoms were constrained with LINCS. ^61^ To remove atomic clashes, the system was energy minimized with the steepest-descent algorithm. We next equilibrated the system under *NVT* conditions for 100 ps at 300 K using the velocity-rescale thermostat^62^ with one heat bath for the coordinated ions, DNA and protein and another heat bath for water and counter-ions. Then, we equilibrated the system at 1 bar for 10 ns under *NPT* conditions using Parrinello-Rahman^63^ pressure coupling and using the same thermostat as in the *NVT* equilibration. During both equilibration steps, all heavy atoms were position restrained with a force constant of 1000 kJ mol^−1^ nm^−2^.

To enable the use a 4 fs time-step for further pulling simulations we used hydrogen mass repartitioning^64^ (HMR). Accordingly, to increase the oscillation period of the bond angles involving hydrogen atoms, the hydrogen masses were scaled up by a factor of *f*_H_ and the heavy atom masses connected to hydrogens were scaled down, while keeping the overall masses of chemical moieties constant. To choose a scaling factor *f*_H_ that yields stable simulations at a 4 fs time step, we tested scaling factors from 2 to 3 in steps of 0.2, where three simulations were carried out for each scaling factor. Each simulation was carried out for 20 ns in *NPT* conditions. None of the simulations with a scaling factor of 2.8 or 3 were stable, whereas all other simulations were stable. For production simulations, we decided to use *f*_H_ = 2.5.

To exclude that HMR leads to excessive energy drift, we carried out three *NVE* simulations with *f*_H_ = 2.5 using 4 fs time step for 500 ps and, for reference, three *NVE* simulation without HMR using a 2 fs time step. On average, we obtained an energy drift of 0.06% ns^−1^ with HMR, which was even smaller than the average value of −0.13% ns^−1^ without HMR (Table S1). Hence, integrating Newton’s equations of motion with HMR models was numerically stable and exhibited only a marginal energy drift.

### Simulation of initial DNA opening pathway

We generated an initial path from the CC to the OC with a steered MD simulation of 175 ns using a combination of one rotational CV and two RMSD-based CVs. As the first CV, we used a rotational CV defined as 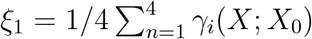, where each dihedral angle *γ*_*i*_ was defined as:

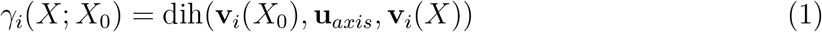

Here, **u**_axis_ denotes the helix axis of the DNA in region +3 to +23 (Fig. S2A). *X*_0_ is the configuration at *t* = 0 ns, and *X* is the configuration at a later simulation time *t*. Further, **v**_*i*_(*X*_0_) denotes the vector connecting the two centers of mass (COMs) **c**_*i*,1_(*X*_0_) and **c**_*i*,2_(*X*_0_) at *t* = 0 ns (Fig. S2B), and **v**_*i*_(*X*) is the instantaneous vector connecting the same COMs at simulation time *t* (Fig. S1C). The two groups of atoms used to define **c**_*i*,1_ and **c**_*i*,2_, respectively, were constructed by splitting the helix DNA region +3 to +23 along the axis, as illustrated in Figs. S2B and S2C. The four **v**_*i*_(*X*_0_) defining *ξ*_1_ are depicted in Fig. S2D.

As a second CV, we used *ξ*_2_ = Δ(*X, X*_OC1_). Here, *X*_OC1_ denotes the DNA backbone atoms of the OC in the region −17 to −5, taken from the 5IYB structure, ^9^ and Δ(*X, X*_OC1_) denotes the RMSD of the instantaneous structure *X* relative to the reference structure *X*_OC1_. As a third CV, we used *ξ*_3_ = Δ(*X, X*_OC2_). Here, *X*_OC2_ denotes the backbone atoms of the OC in the region −17 to +2, again taken from the 5IYB structure.

During the 175 ns of steered MD simulation, we applied different forces on the three CVs described above:

1. The rotational CV (*ξ*_1_) was pulled from 6.27 rad (close to 2*π*) to 0.01 rad over the first 100 ns using a force constant of 7000 kJ mol^−1^ rad^−2^. Over the next 4 ns, the force applied on *ξ*_1_ was turned off by linearly decreasing the force constant from 7000 kJ mol^−1^ rad^−2^ to 0 kJ mol^−1^ rad^−2^.
2. The RMSD relative to *X*_OC1_ (*ξ*_2_) was pulled from 2.35 nm to 0.4 nm over the first 50 ns using a force constant of 10,000 kJ mol^−1^ nm^−2^. Over the next 50 ns, *ξ*_2_ was pulled from 0.4 nm to 0 nm using a force constant decreasing linearly from 10,000 kJ mol^−1^ nm^−2^ to 0 kJ mol^−1^ nm^−2^.
3. The RMSD relative to *X*_OC2_ (*ξ*_3_) was pulled from 1.99 nm to 0 nm between simulation times of 50 ns and 100 ns, using a linearly increasing force constant between 20,000 kJ mol^−1^ nm^−2^ and 30,000 kJ mol^−1^ nm^−2^. Over the next 75 ns, *ξ*_3_ was restrained at 0 nm using a force constant of 30,000 kJ mol^−1^ nm^−2^.

### Relaxation of the initial DNA opening pathway

In order to relax and sample intermediate states along the opening pathway, we first used constant-velocity pulling from the CC to the OC with a Path Collective Variable ^28^ (PCV).

The two components of a PCV are *S*_path_ and *Z*_path_ are defined as:

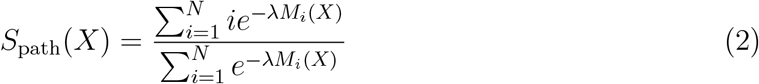

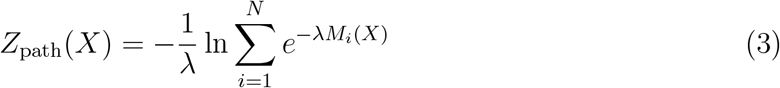

where the unitless *S*_path_ describes the progression along the path and *Z*_path_ describes the deviation from the path. *N* denotes to the number of reference configurations defining the DNA opening path. The distance metric *M*_*i*_(*X*) = Δ^2^(*X, X*_*i*_) is the mean squared deviation (MSD) of the instantaneous configuration *X* relative to the reference configuration *X*_*i*_. The choice of the *N* reference configurations was optimized to obtain similar MSDs between neighboring configurations and a good flatness of the surface spanned by the *N* × *N* MSD matrix.^28,65,66^ The symbol *λ* is the smoothing parameter, proportional to the inverse of the MSD between adjacent reference configurations.

To relax the initial path along *Z*_path_, we performed two rounds of constant-velocity pulling along *S*_path_, while applying a wall potential on *Z*_path_ acting above *Z*_path_(*X*) = 0.035 nm^−2^. For the first relaxation round, we took *N* = 72 reference configurations from the initial path, set *λ* = 21.7 nm^−2^, and we carried out 100 ns of constant-velocity pulling along *S*_path_ from 1.1 to 71.3, corresponding to configurations close to the first or last reference configuration. We used a force constant of 5000 kJ mol^−1^ for *S*_path_ and a force constant of 2.8 × 10^6^ kJ mol^−1^ for *Z*_path_. To build a new path for the following relaxation round satisfying the two criteria for reference selection mentioned above without discarding configurations in the OC, another 20 ns simulation was required with an harmonic restraint centered on *S*_path_(*X*) = 71.3, with a force constant of 5000 kJ mol^−1^ and keeping the wall potential acting above *Z*_path_(*X*) =0.035 nm^2^ with an offset of 0.005 nm^2^ and a force constant of 2.8×10^6^ kJ mol^−1^ nm^−4^. For the second relaxation round, we used *N* = 91 reference configurations from the first relaxation round, set *λ* = 45.43 nm^−2^, and we carried out 100 ns of constant-velocity pulling along *S*_path_ from 1.15 to 90.85 using a force constant of 4000 kJ mol^−1^ for *S*_path_ and a force constant of 4×10^6^ kJ mol^−1^ for *Z*_path_. For the same reasons as for the first relaxation round, we extended the second relaxation simulation for another 20 ns with an harmonic restraint centered on *S*_path_(*X*) = 90.85, with a force constant of 4000 kJ mol^−1^ and keeping the wall potential acting above *Z*_path_(*X*) = 0.035 nm^2^ with an offset of 0.005 nm^2^ and a force constant of 4 × 10^6^ kJ mol^−1^ nm^−4^.

### Sampling the final DNA opening path

From the second relaxation round of constant-velocity pulling described above, we selected *N* = 63 reference configurations to define our final PCV. This final PCV was set up with *λ* = 36 nm^−2^. To sample DNA and loop conformations along the DNA opening path, we extracted 90 configurations from the second round of constant-velocity pulling with *S*_path_ ranging from 1.250 to 62.9. Each of those 90 configurations was used to start a simulation of 50 ns restrained on a particular value of *S*_path_ with an harmonic potential. The reference positions and the force constants of the harmonic potentials of these 90 simulations are shown in Fig. S2. These 90 simulations were used to characterize the opening path in terms of H-bonds, potential energies, and atomic contacts (Figs. 4A and 5).

### Simulation analysis

Hydrogen bonds were defined with a cutoff distance of 0.35 nm between the hydrogen atom and the H-bond acceptor, and with a cutoff angle hydrogen–donor–acceptor of 30°. The base pairs +2 and +3 exhibited disrupted hydrogen bonds but were mismatched with other bases in the downstream DNA fork; hence, during the analysis, we did not consider these bases as part of the open DNA bubble. Contacts were defined with a cutoff distance of 0.3nm. The potential energies were computed as the average of the sum of Lennard-Jones and short-range Coulomb interactions with a cutoff at 1 nm. Simulation trajectories were visualized with PyMOL^67^ and VMD.^68^ Images were generated with PyMOL.

### Conformational stability of the OC

To test the stability of the OC obtained from the second relaxation round, we performed a free simulation of the OC, i.e., without any biasing potential. To this end, a 200 ns simulation was started from the final configuration of the second relaxation round. We quantified the stability of the OC by monitoring the distances between the 12 disrupted base pairs of the transcription bubble (Fig. S3). The 12 distances were computed with the center of geometries of the heavy backbone atoms of the two complementary nucleotides. The 12 distances were then averaged in each time frame. For reference, the same protocol was used to compute the average distances between the 13 dissociated base pairs in the reference OC (5IYB^9^).

### Multiple sequence alignments

RPB1 and RPB2 protein sequences were taken from the UniProtKB/Swiss-Prot ^69^ database. The organisms were chosen to cover different kingdoms of the eukaryotic domain. The alignments were carried out with CLUSTAL W ^35^ version 2.1.

## Acknowledgment

This study was supported by the Deutsche Forschungsgemeinschaft via SFB 860/A16. We thank Christian Dienemann, Sandra Schilbach, Shintaro Aibara, and Patrick Cramer for insightful discussions.

## References

(1) F, C. Central dogma of molecular biology. Nature 1970, 227, 561–563.

(2) He, Y.; Fang, J.; Taatjes, D. J.; Nogales, E. Structural visualization of key steps in human transcription initiation. Nature 2013, 495, 481–486.

(3) Cramer, P.; Bushnell, D. A.; Kornberg, R. D. Structural basis of transcription: RNA polymerase II at 2.8 ångstrom resolution. Science 2001, 292, 1863–1876.

(4) Gnatt, A. L.; Cramer, P.; Fu, J.; Bushnell, D. A.; Kornberg, R. D. Structural basis of transcription: An RNA polymerase II elongation complex at 3.3 Å resolution. Science 2001, 292, 1876–1882.

(5) Nogales, E.; Louder, R. K.; He, Y. Structural Insights into the Eukaryotic Transcription Initiation Machinery. Annual Review of Biophysics 2017, 46, 59–83.

(6) Osman, S.; Cramer, P. Structural Biology of RNA Polymerase II Transcription: 20 Years on. Annual Review of Cell and Developmental Biology 2020, 36, 1–34.

(7) Murakami, K.; Tsai, K. L.; Kalisman, N.; Bushnell, D. A.; Asturias, F. J.; Kornberg, R. D. Structure of an RNA polymerase II preinitiation complex. Proceedings of the National Academy of Sciences of the United States of America 2015, 112, 13543–13548.

(8) Plaschka, C.; Larivière, L.; Wenzeck, L.; Seizl, M.; Hemann, M.; Tegunov, D.; Petrotchenko, E. V.; Borchers, C. H.; Baumeister, W.; Herzog, F.; Villa, E.; Cramer, P. Architecture of the RNA polymerase II-Mediator core initiation complex. Nature 2015, 518, 376–380.

(9) He, Y.; Yan, C.; Fang, J.; Inouye, C.; Tjian, R.; Ivanov, I.; Nogales, E. Near-atomic resolution visualization of human transcription promoter opening. Nature 2016, 533, 359–365.

(10) Plaschka, C.; Hantsche, M.; Dienemann, C.; Burzinski, C.; Plitzko, J.; Cramer, P. Transcription initiation complex structures elucidate DNA opening. Nature 2016, 533, 353–358.

(11) Schilbach, S.; Hantsche, M.; Tegunov, D.; Dienemann, C.; Wigge, C.; Urlaub, H.; Cramer, P. Structures of transcription pre-initiation complex with TFIIH and Mediator. Nature 2017, 551, 204–209.

(12) Dienemann, C.; Schwalb, B.; Schilbach, S.; Cramer, P. Promoter Distortion and Opening in the RNA Polymerase II Cleft. Molecular Cell 2019, 73, 97–106.e4.

(13) Yan, C.; Dodd, T.; He, Y.; Tainer, J. A.; Tsutakawa, S. E.; Ivanov, I.; Ring, M. A. T. Dynamics Provide Insight Into Genetic Diseases. 2019, 26, 397–406.

(14) Schilbach, S.; Aibara, S.; Dienemann, C.; Grabbe, F.; Cramer, P. Structure of RNA polymerase II pre-initiation complex at 2.9 Å defines initial DNA opening. Cell 2021, 184, 4064–4072.e28.

(15) Aibara, S.; Schilbach, S.; Cramer, P. Structures of mammalian RNA polymerase II pre-initiation complexes. Nature 2021, 594, 124–128.

(16) Chen, J.; Chiu, C.; Gopalkrishnan, S.; Chen, A. Y.; Olinares, P. D. B.; Saecker, R. M.; Winkelman, J. T.; Maloney, M. F.; Chait, B. T.; Ross, W.; Gourse, R. L.; Campbell, E. A.; Darst, S. A. Stepwise Promoter Melting by Bacterial RNA Polymerase. Molecular Cell 2020, 78, 275–288.e6.

(17) Huang, X.; Wang, D.; Weiss, D. R.; Bushnell, D. A.; Kornberg, R. D.; Levitt, M. RNA polymerase II trigger loop residues stabilize and position the incoming nucleotide triphosphate in transcription. Proceedings of the National Academy of Sciences of the United States of America 2010, 107, 15745–15750.

(18) Da, L.-T.; Wang, D.; Huang, X. Dynamics of Pyrophosphate Ion Release and Its Coupled Trigger Loop Motion from Closed to Open State in RNA Polymerase II. J. Am. Chem.Soc 2012, 134, 11.

(19) Yu, J.; Bai, L.; Wang, M. D.; Song, Y.-S.; Shu, Y.-G.; Zhou, X.; Da, L.-T.; Huang, X. Constructing kinetic models to elucidate structural dynamics of a complete RNA polymerase II elongation cycle. Physical Biology 2014, 12, 016004.

(20) Silva, D. A.; Weiss, D. R.; Avila, F. P.; Da, L. T.; Levitt, M.; Wang, D.; Huang, X. Millisecond dynamics of RNA polymerase II translocation at atomic resolution. Proceedings of the National Academy of Sciences of the United States of America 2014, 111, 7665–7670.

(21) Zhang, L.; Silva, D. A.; Pardo-Avila, F.; Wang, D.; Huang, X. Structural Model of RNA Polymerase II Elongation Complex with Complete Transcription Bubble Reveals NTP Entry Routes. PLoS computational biology 2015, 11.

(22) Unarta, I. C.; Zhu, L.; Tse, C. K. M.; Cheung, P. P. H.; Yu, J.; Huang, X. Molecular mechanisms of RNA polymerase II transcription elongation elucidated by kinetic network models. Current Opinion in Structural Biology 2018, 49, 54–62.

(23) Tse, C. K. M.; Xu, J.; Xu, L.; Sheong, F. K.; Wang, S.; Chow, H. Y.; Gao, X.; Li, X.; Cheung, P. P. H.; Wang, D.; Zhang, Y.; Huang, X. Intrinsic cleavage of RNA polymerase II adopts a nucleobase-independent mechanism assisted by transcript phosphate. Nature Catalysis 2019 2:3 2019, 2, 228–235.

(24) Christy Unarta, I.; Cao, S.; Kubo, S.; Wang, W.; Pak-Hang Cheung, P.; Gao, X.; Takada, S.; Huang, X. Role of bacterial RNA polymerase gate opening dynamics in DNA loading and antibiotics inhibition elucidated by quasi-Markov State Model.

(25) Shino, G.; Takada, S. Modeling DNA Opening in the Eukaryotic Transcription Initiation Complexes via Coarse-Grained Models. Frontiers in Molecular Biosciences 2021, 8, 1–12.

(26) Grubm ller, H.; Heymann, B.; Tavan, P. Ligand Binding: Molecular Mechanics Calculation of the Streptavidin-Biotin Rupture Force. Science 1996, 271, 997–999.

(27) Leech, J.; Prins, J. F.; Hermans, J. SMD: Visual steering of molecular dynamics for protein design. IEEE computational science engineering 1996, 3, 38–45.

(28) Branduardi, D.; Gervasio, F. L.; Parrinello, M. From A to B in free energy space. The Journal of Chemical Physics 2007, 126, 054103.

(29) Pal, M.; Ponticelli, A. S.; Luse, D. S. The Role of the Transcription Bubble and TFIIB in Promoter Clearance by RNA Polymerase II. Molecular Cell 2005, 19, 101–110.

(30) Lankaš, F.; Lavery, R.; Maddocks, J. H. Kinking Occurs during Molecular Dynamics Simulations of Small DNA Minicircles. Structure 2006, 14, 1527–1534.

(31) Randall, G. L.; Zechiedrich, L.; Pettitt, B. M. In the absence of writhe, DNA relieves torsional stress with localized, sequence-dependent structural failure to preserve B-form. Nucleic Acids Research 2009, 37, 5568–5577.

(32) Mitchell, J. S.; Laughton, C. A.; Harris, S. A. Atomistic simulations reveal bubbles, kinks and wrinkles in supercoiled DNA. Nucleic Acids Research 2011, 39, 3928–3938.

(33) Irobalieva, R. N.; Fogg, J. M.; Catanese, D. J.; Sutthibutpong, T.; Chen, M.; Barker, A. K.; Ludtke, S. J.; Harris, S. A.; Schmid, M. F.; Chiu, W.; Zechiedrich, L. Structural diversity of supercoiled DNA. Nature Communications 2015, 6.

(34) Parvin, J. D.; Sharp, P. A. DNA topology and a minimal set of basal factors for transcription by RNA polymerase II. Cell 1993, 73, 533–540.

(35) Thompson, J. D.; Higgins+, D. G.; Gibson, T. J. CLUSTAL W: improving the sensitivity of progressive multiple sequence alignment through sequence weighting, position-specific gap penalties and weight matrix choice. Nucleic Acids Research 1994, 22, 4673–4680.

(36) Feklistov, A.; Darst, S. A. Structural basis for promoter -10 element recognition by the bacterial RNA polymerase σ subunit. Cell 2011, 147, 1257–1269.

(37) Klimasauskas, S.; Kumar, S.; Roberts, R. J.; Cheng, X. Hhal methyltransferase flips its target base out of the DNA helix. Cell 1994, 76, 357–369.

(38) Huang, N.; Banavali, N. K.; MacKerell, A. D. Protein-facilitated base flipping in DNA by cytosine-5-methyltransferase. Proceedings of the National Academy of Sciences of the United States of America 2003, 100, 68–73.

(39) Naji, S.; Bertero, M. G.; Spitalny, P.; Cramer, P.; Thomm, M. Structure-function analysis of the RNA polymerase cleft loops elucidates initial transcription, DNA unwinding and RNA displacement. Nucleic Acids Research 2008, 36, 676–687.

(40) Barnes, C. O.; Calero, M.; Malik, I.; Graham, B. W.; Spahr, H.; Lin, G.; Cohen, A. E.; Brown, I. S.; Zhang, Q.; Pullara, F.; Trakselis, M. A.; Kaplan, C. D.; Calero, G. Crystal Structure of a Transcribing RNA Polymerase II Complex Reveals a Complete Transcription Bubble. Molecular Cell 2015, 59, 258–269.

(41) Torrie, G. M.; Valleau, J. P. Monte Carlo free energy estimates using non-Boltzmann sampling: Application to the sub-critical Lennard-Jones fluid. Chemical Physics Letters 1974, 28, 578–581.

(42) Laio, A.; Parrinello, M. Escaping free-energy minima. 2002,

(43) Fukunishi, H.; Watanabe, O.; Takada, S. On the Hamiltonian replica exchange method for efficient sampling of biomolecular systems: Application to protein structure prediction. The Journal of Chemical Physics 2002, 116, 9058.

(44) Shaw, D. E. et al. Anton 3: Twenty Microseconds of Molecular Dynamics Simulation before Lunch. International Conference for High Performance Computing, Networking, Storage and Analysis, SC 2021,

(45) Holstege, F. C.; Fiedler, U.; Timmers, H. T. M. Three transitions in the RNA polymerase II transcription complex during initiation. EMBO Journal 1997, 16, 7468–7480.

(46) Kim, T. K.; Ebright, R. H.; Reinberg, D. Mechanism of ATP-dependent promoter melting by transcription factor IIH. Science 2000, 288, 1418–1421.

(47) Gru‥nberg, S.; Warfield, L.; Hahn, S. Architecture of the RNA polymerase II preinitiation complex and mechanism of ATP-dependent promoter opening. Nature Structural Molecular Biology 2012 19:8 2012, 19, 788–796.

(48) Fishburn, J.; Tomko, E.; Galburt, E.; Hahn, S. Double-stranded DNA translocase activity of transcription factor TFIIH and the mechanism of RNA polymerase II open complex formation. Proceedings of the National Academy of Sciences of the United States of America 2015, 112, 3961–3966.

(49) Alekseev, S.; Nagy, Z.; Sandoz, J.; Weiss, A.; Egly, J. M.; Le May, N.; Coin, F. Transcription without XPB Establishes a Unified Helicase-Independent Mechanism of Promoter Opening in Eukaryotic Gene Expression. Molecular Cell 2017, 65, 504–514.e4.

(50) Kostrewa, D.; Zeller, M. E.; Armache, K. J.; Seizl, M.; Leike, K.; Thomm, M.; Cramer, P. RNA polymerase II-TFIIB structure and mechanism of transcription initiation. Nature 2009, 462, 323–330.

(51) Abraham, M. J.; Murtola, T.; Schulz, R.; Páll, S.; Smith, J. C.; Hess, B.; Lindah, E. GROMACS: High performance molecular simulations through multi-level parallelism from laptops to supercomputers. SoftwareX 2015, 1-2, 19–25.

(52) Tribello, G. A.; Bonomi, M.; Branduardi, D.; Camilloni, C.; Bussi, G. PLUMED 2: New feathers for an old bird. Computer Physics Communications 2014, 185, 604–613.

(53) Bonomi, M. et al. Promoting transparency and reproducibility in enhanced molecular simulations. Nature Methods 2019 16:8 2019, 16, 670–673.

(54) E, K.; G, V. YASARA View - molecular graphics for all devices - from smartphones to workstations. Bioinformatics (Oxford, England) 2014, 30, 2981–2982.

(55) Jorgensen, W. L.; Chandrasekhar, J.; Madura, J. D.; Impey, R. W.; Klein, M. L. Comparison of simple potential functions for simulating liquid water. 1983,

(56) Zgarbová, M.; Šponer, J.; Otyepka, M.; Cheatham, T. E.; Galindo-Murillo, R.; Jurečka, P. Refinement of the Sugar-Phosphate Backbone Torsion Beta for AMBER Force Fields Improves the Description of Z-and B-DNA. Journal of Chemical Theory and Computation 2015, 11, 5723–5736.

(57) Maier, J. A.; Martinez, C.; Kasavajhala, K.; Wickstrom, L.; Hauser, K. E.; Simmerling, C. ff14SB: Improving the Accuracy of Protein Side Chain and Backbone Parameters from ff99SB. Journal of Chemical Theory and Computation 2015, 11, 3696–3713.

(58) Macchiagodena, M.; Pagliai, M.; Andreini, C.; Rosato, A.; Procacci, P. Upgrading and Validation of the AMBER Force Field for Histidine and Cysteine Zinc(II)-Binding Residues in Sites with Four Protein Ligands. Journal of Chemical Information and Modeling 2019, 59, 3803–3816.

(59) Darden, T.; York, D.; Pedersen, L. Particle mesh Ewald: An N·log(N) method for Ewald sums in large systems. The Journal of Chemical Physics 1993, 98, 10089–10092.

(60) Miyamoto, S.; Kollman, P. A. Settle: An analytical version of the SHAKE and RATTLE algorithm for rigid water models. Journal of Computational Chemistry 1992, 13, 952–962.

(61) Hess, B.; Bekker, H.; Berendsen, H. J. C.; Fraaije, J. G. E. M. LINCS: A Linear Constraint Solver for Molecular Simulations. J Comput Chem 1997, 18, 14631472.

(62) Bussi, G.; Donadio, D.; Parrinello, M. Canonical sampling through velocity rescaling. The Journal of Chemical Physics 2007, 126, 014101.

(63) Parrinello, M.; Rahman, A. Polymorphic transitions in single crystals: A new molecular dynamics method. Journal of Applied Physics 1998, 52, 7182.

(64) Feenstra, K. A.; Hess, B.; Berendsen, H. J. C. Improving Efficiency of Large Time-Scale Molecular Dynamics Simulations of Hydrogen-Rich Systems. Journal of Computational Chemistry 1999, 20.

(65) Plumed tutorial, Belfast tutorial: Adaptive variables I. https://www.plumed.org/doc-v2.7/user-doc/html/belfast-2.html, Accessed: 2022-01-19.

(66) Plumed tutorial, MARVEL tutorial: Path CVs. https://www.plumed.org/doc-v2.7/user-doc/html/marvel-2.htmluser-doc/html/marvel-2.html, Accessed: 2022-01-19.

(67) Schr‥odinger, L.; DeLano, W. The PyMOL Molecular Graphics System, Version 2.3.0. http://www.pymol.org/pymol.

(68) Humphrey, W.; Dalke, A.; Schulten, K. VMD – Visual Molecular Dynamics. Journal of Molecular Graphics 1996, 14, 33–38.

(69) Bateman, A. et al. UniProt: The universal protein knowledgebase in 2021. Nucleic Acids Research 2021, 49, D480–D489.

