## Supplementary material for "DNA opening during transcription initiation by RNA polymerase II in atomic detail"

### Supporting material for: DNA opening during transcription initiation by RNA polymerase II in atomic detail

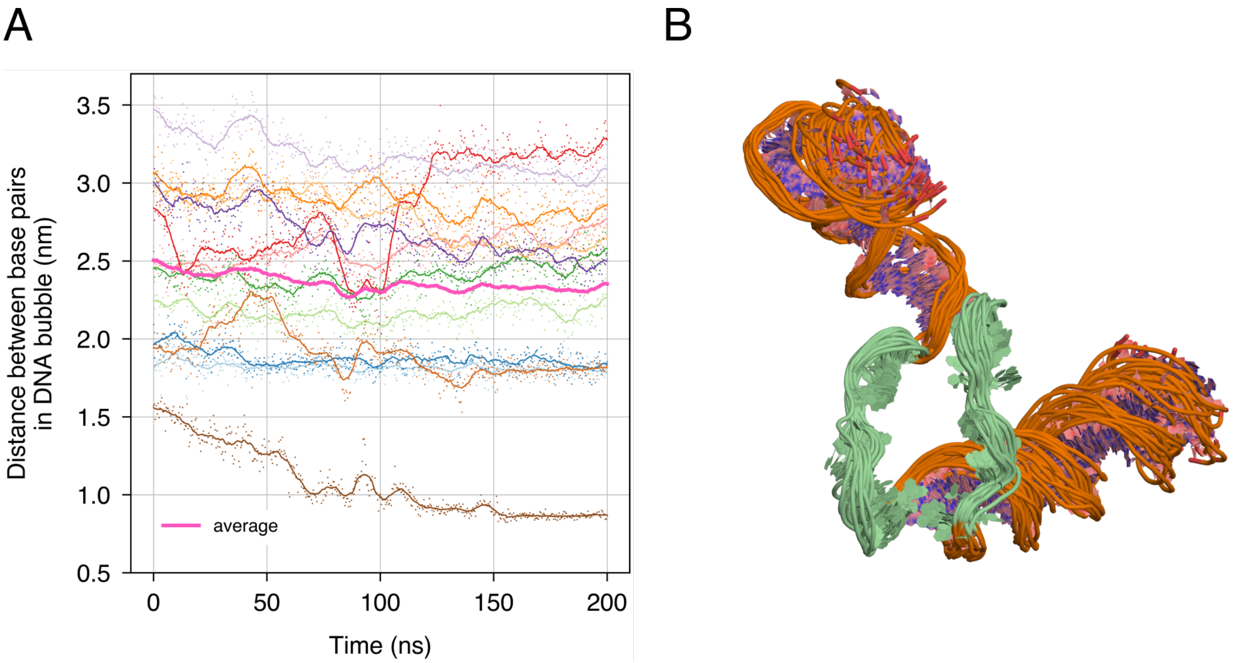


Figure S1: Analysis of the stability of the open DNA bubble in free simulation. (A) To test the stability of the DNA transcription bubble, a free simulation of 200 ns was carried out, starting from the final configuration of the pulling simulation along the final $S_{\text{path}}$. To quantify the stability of the DNA bubble, we monitored the DNA backbone COM distance between disrupted base pairs of the transcription bubble during the free simulation: base pair $-11$, $-10$, $-9$, $-8$, $-7$, $-6$, $-5$, $-4$, $-3$, $-2$, $-1$ and $+1$ are depicted in light blue, dark blue, light green, dark green, light pink, red, light orange, dark orange, light purple, dark purple, and maroon, respectively. The average distance between the 12 base pairs is depicted in dark pink. (B) DNA snapshots taken every 10 ns from our free simulation. Overall, distances between disrupted base pair and visual inspection of the trajectory show that the two strands do not re-anneal, suggesting that a stable DNA bubble was obtained in our simulation.

Table S1: Drifts of the total simulation energy during three independent *NVE* simulations of 500 ps with HMR using a 4 fs time step (left column) or without HMR using a 2 fs time step (right column). Energy drifts are shown relative to the total energy of the simulation per nanosecond. Using HMR does not increase the energy drift.

| Total energy drift with HMR ($\%\,\text{ns}^{-1}$) | Total energy drift without HMR ($\%\,\text{ns}^{-1}$) |
|---|---|
| 0.06 | -0.12 |
| 0.06 | -0.14 |
| 0.06 | -0.12 |

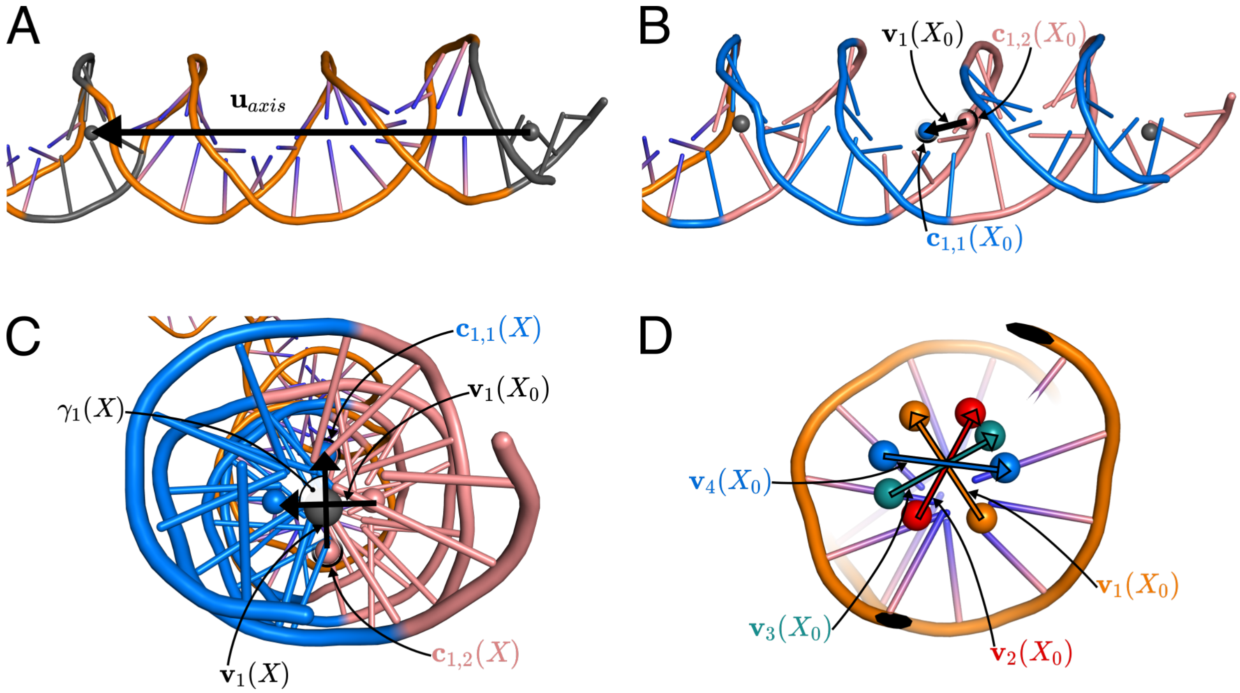


Figure S2: Illustration of the definition of the rotational CV $\xi_1$. (A) Axis of the DNA helix used to define one vector of the dihedral angles in region +3 to +23. Residues colored in grey were used to compute the center of geometries defining the axis (gray beads). (B-C) Pink and blue DNA highlight two horizontal halves of the DNA, the pink bead is located at the center of geometry of the pink half, and the blue bead is located at the center of geometry of the blue half. (D) Colored beads represent the center of geometry of an horizontal half of the DNA region used for the rotation. Vectors $\mathbf{v}_1$ through $\mathbf{v}_4$ connecting pairs of beads with identical colors were used to define the four dihedral angles $\gamma_1$ through $\gamma_4$, see Methods.

| $S_{path}$ | $\kappa$ ($kJ\,mol^{-1}$) |
|---|---|
| 1.250 | 10 |
| 2.244 | 10 |
| 3.239 | 10 |
| 4.233 | 10 |
| 5.227 | 10 |
| 6.222 | 10 |
| 7.216 | 10 |
| 8.210 | 10 |
| 9.205 | 10 |
| 10.199 | 10 |
| 11.194 | 10 |
| 12.188 | 10 |
| 13.182 | 10 |
| 14.177 | 10 |
| 15.171 | 10 |
| 16.165 | 10 |
| 17.160 | 10 |
| 18.154 | 10 |

| $S_{path}$ | $\kappa$ ($kJ\,mol^{-1}$) |
|---|---|
| 19.148 | 10 |
| 20.143 | 10 |
| 21.137 | 10 |
| 22.131 | 10 |
| 23.126 | 10 |
| 24.120 | 10 |
| 25.115 | 10 |
| 26.109 | 10 |
| 27.103 | 10 |
| 27.200 | 20 |
| 27.900 | 20 |
| 28.098 | 10 |
| 29.092 | 10 |
| 30.086 | 10 |
| 31.081 | 10 |
| 32.075 | 10 |
| 33.069 | 10 |
| 34.064 | 10 |

| $S_{path}$ | $\kappa$ ($kJ\,mol^{-1}$) |
|---|---|
| 34.100 | 20 |
| 34.600 | 20 |
| 35.025 | 20 |
| 35.058 | 10 |
| 35.950 | 20 |
| 36.052 | 10 |
| 36.875 | 20 |
| 37.047 | 10 |
| 37.800 | 20 |
| 38.041 | 10 |
| 39.035 | 10 |
| 40.030 | 10 |
| 41.024 | 10 |
| 42.019 | 10 |
| 43.013 | 10 |
| 44.007 | 10 |
| 45.002 | 10 |
| 45.996 | 10 |

| $S_{path}$ | $\kappa$ ($kJ\,mol^{-1}$) |
|---|---|
| 46.000 | 20 |
| 46.840 | 20 |
| 46.990 | 10 |
| 47.680 | 20 |
| 47.985 | 10 |
| 48.520 | 20 |
| 48.979 | 10 |
| 49.360 | 20 |
| 49.973 | 10 |
| 50.200 | 20 |
| 50.550 | 20 |
| 50.968 | 10 |
| 51.962 | 10 |
| 52.956 | 10 |
| 53.000 | 20 |
| 53.750 | 20 |
| 53.951 | 10 |
| 54.500 | 20 |

| $S_{path}$ | $\kappa$ ($kJ\,mol^{-1}$) |
|---|---|
| 54.945 | 10 |
| 55.250 | 20 |
| 55.940 | 10 |
| 56.000 | 20 |
| 56.750 | 20 |
| 56.934 | 10 |
| 57.500 | 20 |
| 57.928 | 10 |
| 58.923 | 10 |
| 59.917 | 10 |
| 60.000 | 20 |
| 60.911 | 10 |
| 60.933 | 20 |
| 61.867 | 20 |
| 61.906 | 10 |
| 62.300 | 8 |
| 62.800 | 20 |
| 62.900 | 10 |

Figure S3: Tables of harmonic potential centers $S_{\mathrm{path}}$ and force constants $\kappa$ used to sample the DNA opening path.

Movie S1: Overview of the relaxed CC-to-OC transition obtained by a pulling simulation along $S_{\mathrm{path}}$, shown from three different viewpoints. The progress bar indicates the progression along $S_{\mathrm{path}}$. Protein and DNA is shown as cartoon, Zinc ions as black spheres. The rudder, fork loop 1, and fork loop 2 are shown as tubes in fuchsia, beige and cyan, respectively. Colors are consistent with Fig. 1.

Movie S2: Atomic view of the relaxed CC-to-OC transition obtained by pulling simulation along $S_{\mathrm{path}}$, the focus is made on the transcription bubble region. A progress bar indicates the position of each frame along $S_{\mathrm{path}}$. Colors are consistent with Fig. 1.

Movie S3: Fork loop 2 flipping during the relaxed CC-to-OC transition obtained by pulling simulation along $S_{\mathrm{path}}$, shown from two different viewpoints. The progress bar indicates the progression along $S_{\mathrm{path}}$. Colors are consistent with Fig. 1.
